# Differences in chromatin accessibility between renal cortex and inner medulla correlate with spatial differences in gene expression and are modulated by NFAT5 function

**DOI:** 10.1101/2024.04.23.589187

**Authors:** Kristina Engel, Dmitry Chernyakov, Katrin Nerger, Katrin Sameith, Andreas Dahl, Bayram Edemir

## Abstract

A spatial gene expression pattern between the cortex (CTX) and inner medulla (IM) of the kidney has been observed, but the underlying mechanisms are unclear. Understanding these mechanisms is essential for elucidating renal function. Using the Assay for Transposase-Accessible Chromatin with high-throughput sequencing (ATAC-seq) we analyzed the open chromatin structures and the involvement of epigenetic mechanisms in mediating gene expression differences between the renal CTX and IM. We also examined the role of the nuclear factor of activated T cells 5 (NFAT5), a key regulator of hypertonicity. ATAC-seq analysis was performed on CTX and IM samples from both wild-type (WT) and NFAT5 knockout (KO) mice.

This work demonstrates for the first time that these differences in gene expression between renal CTX and IM are associated with an epigenetic mechanism driven by chromatin accessibility, which is partially modulated by the nuclear factor of activated T-cells 5 (NFAT5) in mice. Furthermore, spatial localization and NFAT5-promoted chromatin accessibility correlate with differential gene expression and altered promoter binding motif enrichment in CTX and IM.

This study provides new insights into the spatial and NFAT5-mediated regulation of chromatin accessibility and gene expression in CTX and IM. This work advances our understanding of kidney physiology by uncovering previously unknown epigenetic factors influencing gene expression and provides a new perspective on renal adaptive mechanisms.

**TRANSLATIONAL STATEMENT:** The study reveals new insights into the spatial and epigenetic regulation of gene expression in the renal cortex (CTX) and inner medulla (IM) in the mouse kidney. We used the Assay for Transposase-Accessible Chromatin with High-Throughput Sequence Analysis (ATAC-seq) to identify a key role of NFAT5 in modulating chromatin accessibility and to uncover previously unknown epigenetic factors. This research enhances our understanding of renal physiology and has important implications for clinical care by providing insights into potential adaptive mechanisms in the kidney. These findings suggest future investigations targeting epigenetic signaling pathways for therapeutic intervention in renal diseases.

## INTRODUCTION

The mammalian kidney is a complex organ comprised of individual nephron units, including the glomerulus and renal tubules, each with specific functions related to urine production. These segments contain approximately 20 specialized cell types, each with distinct gene expression profiles ^1^. Recent advancements in sequencing techniques, such as Next Generation Sequencing (NGS) and single cell RNA-sequencing (snRNA-seq), have allowed for the identification of nephron segment-or cell-specific gene expression patterns in the kidney ^2, 3^. For example, snRNA-seq was used to profile the transcriptome of the major collecting duct cell types ^4^. These techniques have been applied to study gene expression profiles under physiological ^5–8^ and pathophysiological conditions in the kidney ^9–11^. The mechanisms that lead to these changes in gene expression and the consequences of these changes on progression or development of the disease state are not fully understood. A recent study, using snRNA-seq, showed that the corticomedullary osmotic gradient in the kidney plays a crucial role in urine concentration and also influences gene expression profiles ^12^. The generation of this gradient relies on the water absorption in the renal collecting ducts (CD) ^13^ and the reabsorption of NaCl in the loop of Henle, leading to the production of highly concentrated urine ^13–16^. This process creates a hypertonic environment, reaching up to 1200 mosmol/kg in humans and 4000 mosmol/kg in mice, within the renal inner medulla ^17^. While hypertonicity can induce DNA damage and cell death in most cells, the cells of the renal medulla have developed adaptive mechanisms to maintain their function and survive They accumulate compatible organic osmolytes, such as taurine, myo-inositol, betaine, or sorbitol, facilitated by specific membrane transporters like myo-inositol transporter (SLC5A3) ^18^, sodium coupled betaine transporter (SLC6A12) ^19, 20^, sodium coupled taurine transporter (SLC6A6) ^20^ or enzymes, like aldose reductase (AKR1B1) ^21^. One of the key regulators in these adaptive processes is the transcription factor NFAT5, also known as tonicity-responsive enhancer-binding factor (TonEBP). NFAT5 is activated under hypertonic conditions and translocates into the nucleus ^22^, where it induces the expression of genes involved in the accumulation of organic osmolytes, as well as heat shock proteins ^23–25^. Additionally, NFAT5 is implicated in the regulation of urea transporters ^26^ and the water channel aquaporin-2 (AQP2) ^27^, These genes were also found to have hypertonicity-induced increase of gene expression ^28, 29^. Despite the established role of NFAT5 in response to hypertonicity, its functions independent of hypertonic challenge and its comprehensive impact on renal gene expression profiles remain poorly understood. Additionally, the factors controlling the spatial gene expression pattern along the corticomedullary axis, associated with the osmotic gradient, are not fully understood. Epigenetic modifications, such as DNA methylation and histone modifications, are known to influence chromatin accessibility and induce changes in gene expression ^30^. Therefore, the analysis of epigenetic modifications under hypertonic conditions or the corticomedullary osmotic gradient could provide insights into the differences observed in gene expression patterns.

Recently, Haug et al. published a study on human kidney samples, using gene expression profiling and epigenetic changes to characterize spatial differences in gene expression between the renal cortex and inner medulla ^31^. As described above one factor that influences spatial gene expression is the corticomedullary osmotic gradient and the action of NFAT5. This research represents a valuable addition to the study of Haug et al. since we also show the impact of NFAT5 on gene expression by epigenetic modifications. The aim of this study is to investigate the chromatin accessibility underlying gene expression in the renal medulla, compared to the renal cortex in mice. We examined chromatin accessibility in the renal cortex and inner medulla of control and PC-deficient NFAT5 mice and correlated these results with RNA-Seq data obtained in a previous study. Through these investigations, the present study is expected to provide valuable insights into NFAT5-promoted chromatin accessibility and gene expression regulation in the kidney, and to highlight the importance of NFAT5 in renal function.

## METHODS

### Mouse Strains

To generate mice (strain C57BL/6N) that are deficient in NFAT5 in the principal cell (PC) of the collecting duct, floxed NFAT5 mice (NFAT5-fl/fl) were crossed with Aqp2-Cre mice expressing CRE recombinase under the control of the Aqp2 gene promoter ^29, 32^. The insertion of the Cre recombinase disrupts one allele of the Aqp2 gene. Therefore, we used only NFAT5fl/fl-AQP2-470 CRE+/− mice with Aqp2-Cre+/− mice as controls. The mice were bred in the research laboratory’s own animal facility. For the ATAC-seq the cortex and the inner medulla tissue of the same kidney were used. In total, kidneys from n = six adult female mice were used, n = 3 mice were from the NFAT5-KO group and n = 3 from the Aqp2-Cre+/− control group. There were no further inclusion or exclusion criteria for either test group. The mice for each group came from one cage and three animals were taken from each group. All mice were were kept in individually ventilated cages (IVCs) at a temperature of 20°C to 24°C, humidity of 45% to 60%, 15 air exchanges per hour. Feeding (1314 breeding diet from Altromin, Lage, Germany) and drinking ad libitum,. The animal experiments in this study were approved by the Commission for Laboratory Animal Husbandry of the State of Saxony-Anhalt, Germany, and conducted according to local guidelines (approval code: 203.h-42502-2-1250 MLU). The NFAT5-fl/fl mice were kindly provided by Prof. C. Küper ^33^. The Aqp2-Cre deleted mice were kindly provided by Dr Juliette Hadchouel ^34^.

### RNA-seq data

The RNA-seq data were collected in a previous study (9) and have been submitted to the public repository Gene Expression Omnibus (GSE195881, https://www.ncbi.nlm.nih.gov/geo/query/acc.cgi?acc=GSE195881).

### Tissue sample preparation for ATAC-seq

Fresh kidneys were removed from the mice. A small piece of renal CTX tissue (approx. 2 mm) and the renal IM were transferred to 5 ml ice-cold PBS (Thermo Scientific). After centrifugation at 500 x g for 5 min at 4°C, the PBS was aspirated and 500 µl ATAC lysis buffer (ActiveMotif ATAC-seq kit, Carlsbad, CA) was added. Samples were homogenized using the gentleMACS™ Octo Dissociator with Heaters (Miltenyi Biotec, Germany) and the gentleMACS program 37C_Multi_E at 37°C for 31 min. After filtering through a 100 µm mesh strainer the strainer was rinsed with 15 ml of PBS. Cells were harvested for 10 min at 300 x and eluted in 5 ml PBS for CTX and 1 ml PBS for IM followed by counting.

### ATAC-seq

ATAC was performed with 100.000 cells per sample (*n* = 3 biological replicates of CTX and IM from three kidneys per group, total n = 12 samples). We chose a small sample size because the NFAT5 knockout was evaluated in vivo using ATAC-seq for the first time in the present study and therefore the initial intention was to collect basic findings. The following parameters were assessed: number of ATAC-seq peaks and number of differential accessible regions (DARs). We used the ActiveMotif ATAC-seq kit (Carlsbad, CA), according to the manufacturer’s instructions (manual version B6) for tagmentation and purification. For the library preparation a total of 10 µl of purified tagmented DNA was indexed and pre-amplified for initial 5 PCR cycles with 1x KAPA HiFi HotStart Readymix and 100 nM unique dual index P5 and P7 primers compatible with Illumina Nextera DNA barcoding, under the following PCR conditions: 72°C for 5 min, 98°C for 30 s, thermocycling for 5 cycles at 98°C for 10 s, 63°C for 30 s and 72°C for 1 min. Subsequently, a qPCR on the LightCycler 480 (Roche) was performed with 1 µl of the pre-amplified material to determine the remaining PCR cycle numbers (8, 9, or 11) to avoid saturation and potential biases in library amplification ^35^. Purification and double-sided size selection of amplified libraries was done with AMPure XP beads (Beckmann Coulter; starting with a 1.55x volume of XP bead purification, followed by a 0.6x/1.55x double-sided size selection), checked for their quality on a Fragment Analyzer (Agilent) and quantified using qPCR. Libraries were sequenced on the Illumina NovaSeq 6000 with PE 100 bp reads to a depth of 60-300 M read pairs.

### ATAC-seq data analysis

ATAC-seq bioinformatics analysis was done using the nf-core atacseq pipeline v1.2.1 ^36^, with “--genome GRCm39--macs_gsize 2468088461”, and default settings otherwise. The pipeline is very comprehensive and automatically runs all required steps on a library-level, including (but not limited to) raw read QC, adapter trimming, alignment, filtering, generation of normalized coverage tracks, peak calling, and peak annotation based on the closest neighboring gene and genomic features (promoter-transcription start site (TSS), exon, transcription termination site (TTS), intron, intergenic). The evaluation is based on the analysis of fragments. The broad peak results of three replicates per tissue were merged. For visualization, bigwig files were generated. Finally, a consensus peak set is created across all libraries, and differential accessibility analysis performed between CTX and IM and between WT and the respective NFAT5-KO using DESeq2 ^37^. The resulting *P* values were adjusted using the Benjamini and Hochberg approach to control for false discovery rate (FDR). Broad peaks with an absolute fold-change ≥ 1 and FDR ≤ 0.05 are considered as differentially accessible regions (DARs).

### Integration of ATAC-Seq and RNA-Seq datasets

Differential expressed genes (DEGs) with differential accessible regions (DARs) were identified by overlapping the list of DEGs (FDR ≤ 0.05) with the list of DARs (FDR ≤ 0.05) for CTX and IM, for CTX-KO and CTX-WT and for IM-KO and IM-WT. The proportion of overlapping gene regions (differentially accessible and expressed) in the total proportion of differentially accessible regions was calculated as a percentage. Since a gene can have multiple accessible regions and it is mainly the promoter that is crucial for transcription, the integration was limited to promoter regions.

### Visualization of RNA and ATAC data

Correlation plot, and PCA were generated using the DESeq2 plotPCA function. The heatmap with ATAC values around the TSS of genes and corresponding profile were plotted with computeMatrix (Galaxy Version 3.5.2) and plotHeatmap Galaxy (Version 3.5.2) from deepTools2 ^38^. For the clustered heatmap, the transcripts per million (TPM) values were calculated from the featureCounts (program included in the nf-core atacseq pipeline) raw data and then the z-score was determined from the TPMs and visualized. For visualization of normalized coverage tracks from RNA-seq and ATAC-seq (bam and bigWig files) pyGenomeTracks (Galaxy Version 3.8+galaxy2, ^39^) was applied including the gencode_vM2_primary_assembly_annotation gtf file. GraphPad Prism 8 software was used to generate volcano plots including the calculation of the Pearson correlation coefficient r to investigate the relationship between gene expression and chromatin accessibility. Pearson correlation coefficient r_sig_ was calculated from significant differentially expressed genes with differentially accessible regions (|log_2_FC| ≥ 1). Pearson correlation r_all_ was calculated from significant (|log_2_FC| ≥ 1) and non-significant (|log_2_FC| ≤ 1) differentially expressed genes with differentially accessible regions.

### Motif enrichment analysis

Motif analysis to infer transcription factor binding was performed using the findMotifsGenome program (Galaxy version 4.11+galaxy3,^40^) from Homer to determine motif enrichment of known motifs in accessible chromatin regions. Known motifs with a q-value ≤ 0.05 were assumed to be significant.

### Gene ontology analysis

Gene Ontology (GO) functional enrichment analyses were performed to annotate genes with differential accessible promoters to biological processes and pathways using ^41^ ShinyGO 0.7. An adjusted *P* value of 0.05 was used as the threshold for significance of GO terms.

### Statistics and reproducibility

The respective *P* values used to determine the significance were described in the text. The numbers of replicates (n =3) are biological replicates of kidneys from three mice each from WT and NFAT5-KO. A sample of CTX and IM was separated from each individual kidney, resulting in a sample size of *n* =3 each of CTX and IM from the WT and the NFAT5-KO mouse. We used the ARRIVE reporting guidelines ^42^.

## RESULTS

### ATAC-Peak distribution indicates global differences in the accessibility of chromatin in the renal cortex and medulla

In our previous transcriptional analysis, our results showed that hypertonicity and NFAT5 are important regulators of cell type and spatial gene expression in the kidney ^28, 29^. To support our hypothesis that the observed gene expression changes between cortex and medulla are due to epigenetic mechanisms, we analyzed the open chromatin structures using Assay for Transposase-Accessible Chromatin with high-throughput sequencing (ATAC-seq). ATAC-seq allows to quantify genome-wide chromatin accessibility of genomic regions ^43^.

We performed ATAC-seq using renal CTX and IM from control (CTX-WT, IM-WT) and NFAT5-KO (CTX-KO, IM-KO) mice. Hierarchical clustering clearly separated the samples separating CTX from IM samples and control from NFAT5-KO samples, indicating minimal variation during tissue preparation (Fig. 1a). The principal component analysis (PCA) confirmed the differences in chromatin accessibility between the groups (Fig. 1b).

**Figure 1:**
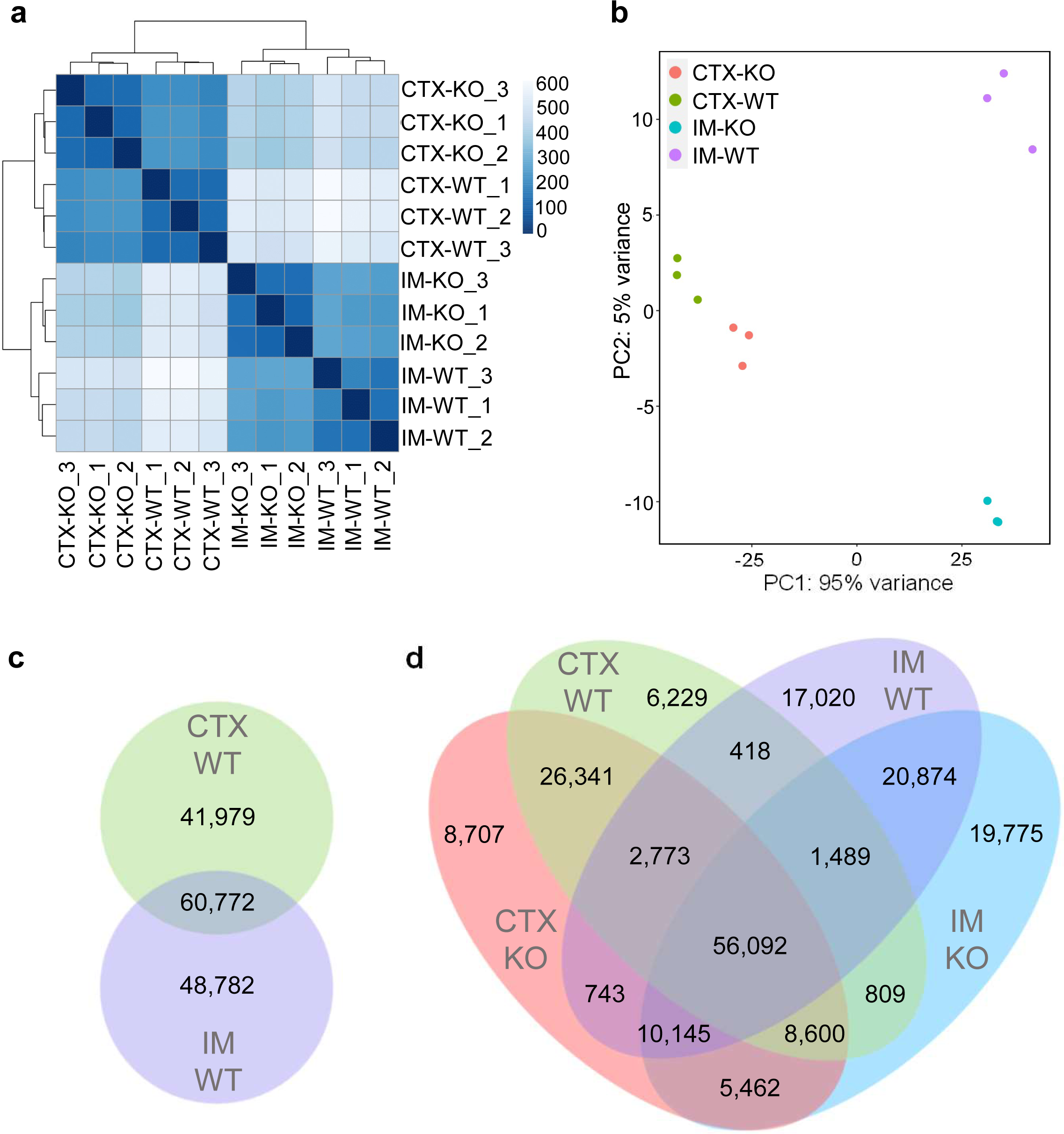
Correlation and intersection analysis in the ATAC-seq samples. (a) Correlation analysis and hierarchical clustering of ATAC-seq peak results from three sample replicates each from CTX and IM of WT and NFAT5-KO mice. The blue color bar represents the correlation efficiency. (b) Principle component analysis of three ATAC-seq sample replicates each from CTX and IM of WT and NFAT5-KO mice. Venn diagram analysis determined overlapping and tissue-specific ATAC-seq intervals between CTX-WT and IM-WT (c) and between CTX-WT, IM-WT, CTX-KO and IM-KO (d).

Table 1 shows the number of ATAC peaks, which represents regions of open chromatin, identified in each of the groups. More than 100,000 different regions were identified. Interestingly, loss of NFAT5 function led to increased number of accessible regions, for example 104,606 in control CTX vs. 120,502 in NFAT5-KO CTX. The list with accessible regions is provided as supplementary data S1.

**Table 1:**
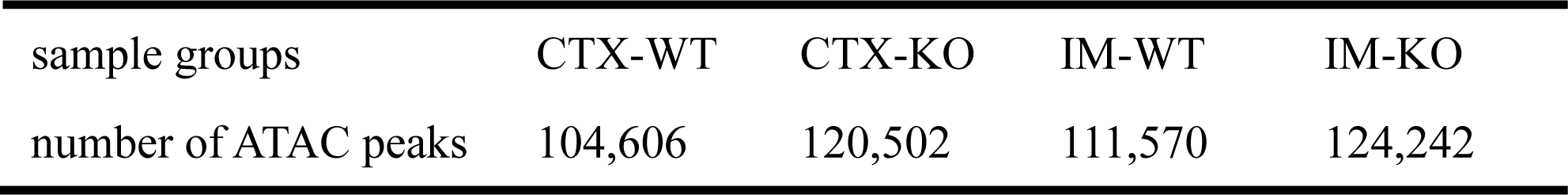
Number of ATAC sequencing peaks in the different groups.

| sample groups | CTX-WT | CTX-KO | IM-WT | IM-KO |
| --- | --- | --- | --- | --- |
| number of ATAC peaks | 104,606 | 120,502 | 111,570 | 124,242 |

In the next step, we compared how many regions are common and unique between CTX and IM in control mice. To perform the comparing analysis, the individual ATAC-seq peak sets identified for each sample were merged to create a consensus set of peaks (supplementary data S2). Using these consensus peaks, we evaluated the amount of overlapping regions between the peaks of a group of samples. In the WT mouse kidney, a total of 151,533 regions with open chromatin were detected, consisting of 41,979 that were unique to CTX, 48,782 unique to IM, and 60,772 regions that were common (Fig. 1c). The loss of NFAT5 was associated with 33,944 additional open chromatin regions in the kidney, with 8,707 being unique to CTX, 19,775 unique to the IM, and 5,462 common regions (Fig. 1d). A total of 56,092 open regions were common across all samples. Moreover, a variety of chromatin regions has been discovered, which are exclusive to specific tissue types and remain unaffected by NFAT5 function. For example, within the IM-WT group, 17,020 regions are unique accessible in this tissue type and are not influenced by NFAT5, while the CTX-WT group exhibits 6,229 regions that are exclusively accessible in that tissue type (Fig. 1d).

The distribution of chromatin accessibility for different genes at their transcription start site (TSS) can be visualized in the different samples. This pattern is indicated by the intensity of the ATAC signal (Fig. 2a, top figure) and the distribution of peaks (Fig. 2a, upper figure), both showing a concentration of signal strength around the promoter-TSS region in all samples. This observation underscores the good quality of the ATAC-seq data and demonstrates the importance of the accessible chromatin regions, which are predominantly located around the TSSs ^44^. These accessible promoter regions are critical for transcription factor binding and transcriptional regulation. We annotated all of the accessible regions and distinguishing them based on their positions within intron, exon, intergenic, promoter-TSS and transcription termination site (TTS). When analyzing the proportions of annotated globally accessible regions, no major differences were found between the different sample types in the first survey. (Fig. 2b). We plotted the TPM values for each peak and created a clustered heatmap with the calculated z-scores to visualize the general differences in accessible promoter regions of protein-coding genes between CTX and IM and WT and KO (Fig. 2c). In general, it is possible to identify promoter regions of closed chromatin in CTX and more accessible chromatin in IM, and vice versa. It is also possible to identify promoter regions within CTX and IM where the loss of NFAT5 leads to a gain or loss of chromatin accessibility. This figure highlights the major differences in chromatin accessibility associated with tissue type and genetic background.

**Figure 2:**
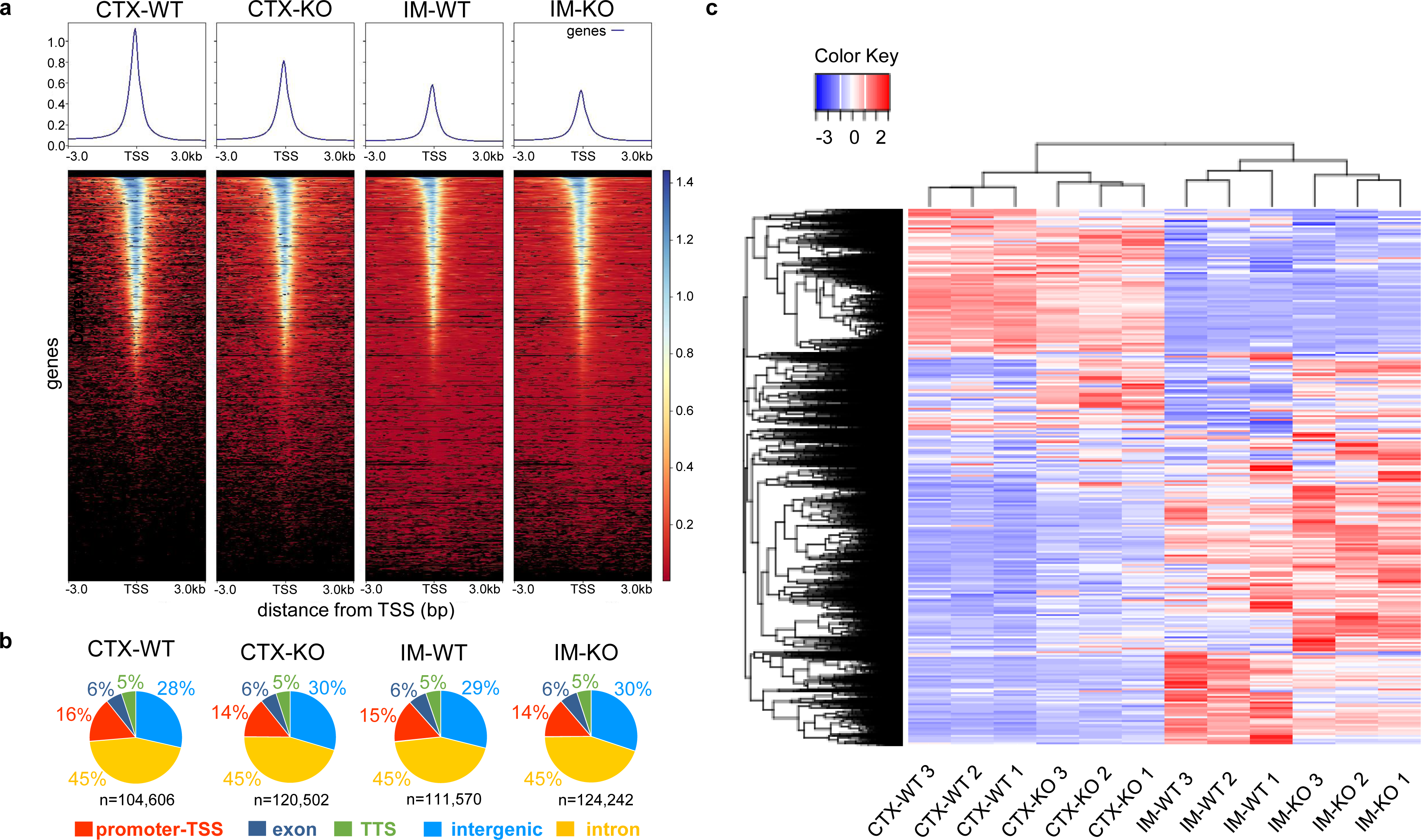
ATAC-seq reveals global differences in the chromatin landscape in the kidney of CTX and IM. (a) Upper, averaged profiles of ATAC-seq signal intensity at the transcription start site (TSS). Bottom, heatmap plots of the signal distribution of ATAC-seq peaks at the TSS. (b) Pie charts show the distribution of genomic features among ATAC-seq peaks. (c) Clustered heatmap comparing z-scores calculated for each row. Each row represents a detected ATAC-seq peak at the promoter-TSS region of accessible protein-coding genes in three replicates of WT and NFAT5-KO from CTX and from IM. Red indicates a more-accessible region and blue indicates a less-accessible region.

### Spatial localization and NFAT5 function mainly affect the differential accessibility of promoters in CTX and IM

To determine the specific differences (gain or loss) in accessibility between CTX and IM and between WT and NFAT5-KO mice, we identified the significantly differential accessible regions (DARs) between the groups. The results are visualized by Volcano plots (Fig. 3a). While 61,846 regions (55 %) show a significant loss of accessibility, 50,782 (45%) regions have a gain of accessibility in CTX-WT compared to IM-WT. The number of DARs is similar to the number of unique accessible regions identified in figure 1c. Between control CTX and NFAT5-KO CTX nearly 19,000 DARs were identified and more than 31,000 DARs between control and NFAT5-KO IM (Fig. 3b).

**Figure 3:**
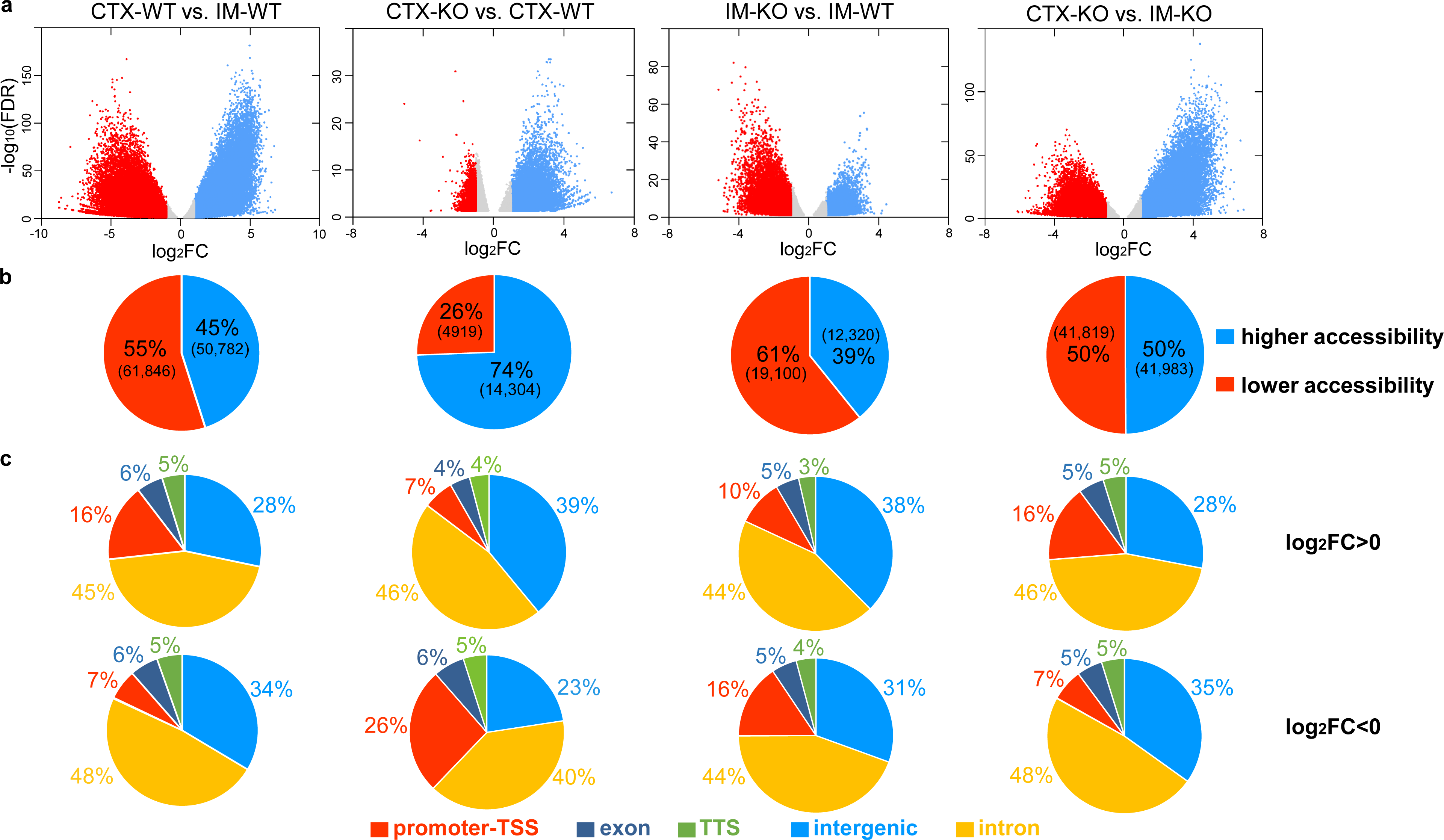
Differentially accessible regions in the CTX and IM are influenced by spatial localization and NFAT5. (a) Volcano Plots of differentially accessible regions (FDR ≤ 0.05; red dots = log_2_FC ≤ −1, blue dots = log_2_FC ≥ 1, grey dots = |log_2_FC| ≤ 1) in WT and NFAT5 knockout tissue of CTX and IM. (b) Visualization of the proportion of differentially accessible regions shows regions in blue that are becoming more accessible and regions in red that are showing a decrease in accessibility (|log_2_FC| ≥ 1; FDR ≤ 0.05). (c) Pie charts show the distribution of genomic features between regions that are becoming more accessible (top, log_2_FC ≥ 0, FDR ≤ 0.05) and regions that are losing their accessibility (bottom, log_2_FC ≤ 0, FDR ≤ 0.05).

The annotation of the differentially accessible regions provides initial evidence of differences in the accessibility between the groups (Fig. 3c). In the CTX-WT, 16 % of DARs with gain of accessibility are localized in promoter regions compared to IM-WT, while only 7 % of DARs regions with loss of accessibility are localized at the promotor regions. The loss of NFAT5 on accessibility within promoter regions was associated with more reduced accessibility in the promotor regions in both CTX and IM. A list with the differentially accessible regions is provided as supplementary data S3.

### The chromatin accessibility at promoters is correlated with RNA expression

A crucial role in the initiation and regulation of the transcription process is assigned to the promoter. To investigate whether the changes in open chromatin regions (DARs) at promoter-TSS correlate with the changes in gene expression (differentially expressed genes = DEGs), we performed an intersection analysis of the ATAC-seq and RNA-seq data set ^29^. We used the RNA-seq data generated from a previous study using CTX and IM from the same control and NFAT5-KO mice used here for ATAC analysis ^29^. For the intersection the lists of differentially up-regulated genes and differentially down-regulated genes were intersected with the list of DARs at the promoter (TSS) regions (supplementary data S4). As a result, four combinations of the regulatory state of gene expression and promoter accessibility are possible: 1. mRNA up /gain of accessibility, 2. mRNA up/loss of accessibility, 3. mRNA down/gain of accessibility and 4. mRNA down/loss of accessibility. Figure 4a shows the results of this comparison. For example, for around 20 % of the DEGs that are up regulated in IM vs. CTX we had a correlation with gain of accessibility and for around 40 % of DEGs that are down regulated in expression, we observed a loss of accessibility.

**Figure 4:**
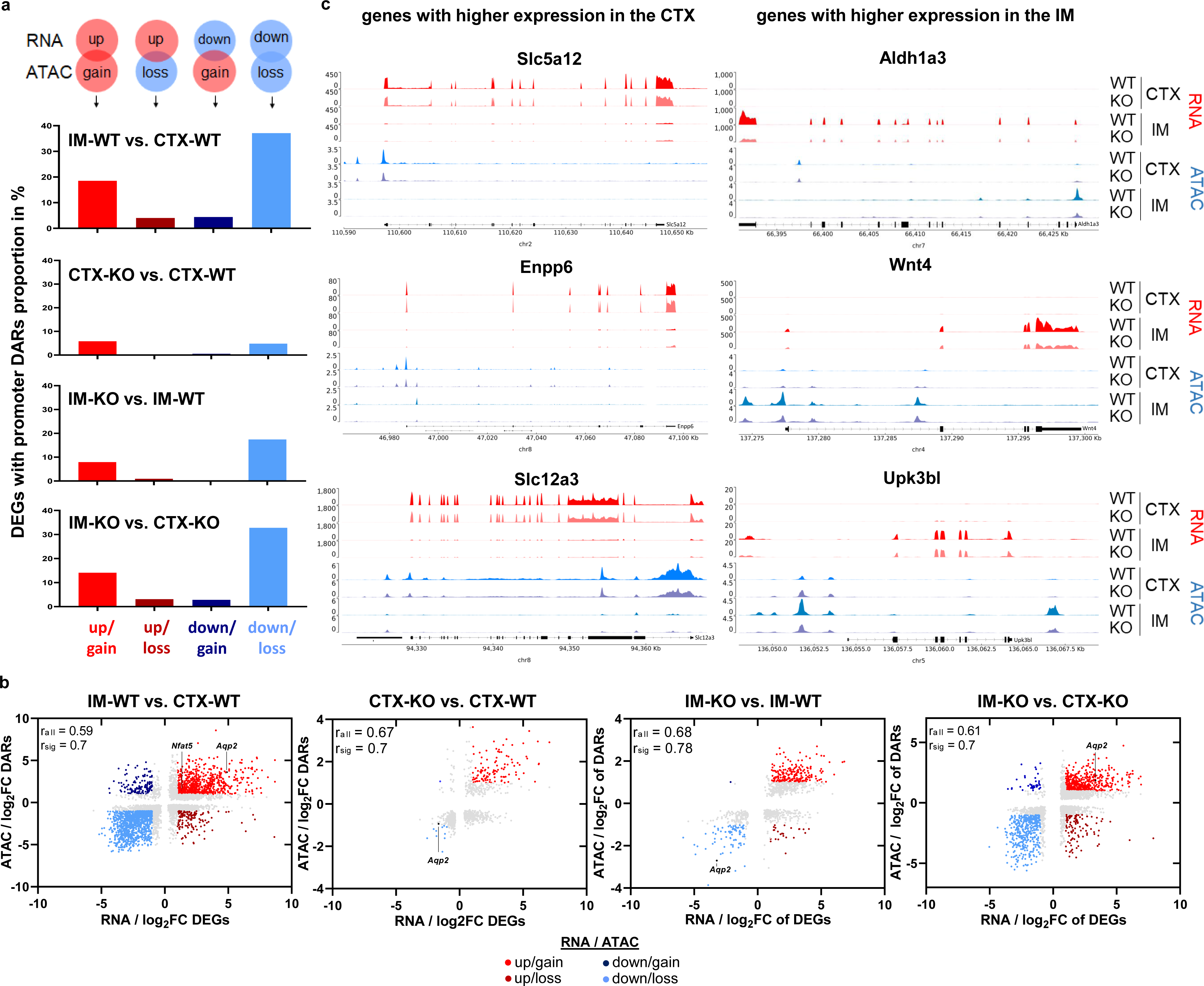
RNA expression is correlated with chromatin accessibility at promoter sites. (a) Comparison of differentially expressed genes (DEGs; FDR ≤ 0.05, |log_2_FC| ≥ 1) with differentially accessible promoter-transcription start site (TSS) region (promoter DARs; FDR ≤ 0.05, |log_2_FC| ≥ 1). Shown is the proportion of genes with up-regulated gene expression and gain-or loss of chromatin accessibility (up/gain and up/loss) and with down-regulated gene expression and gain-or loss of chromatin accessibility (down/gain and down/loss) in the four gene sets of the CTX and IM of WT and NFAT5-KO mice. (b) Correlation of log_2_FC of DEGs (FDR ≤ 0.05) of IM vs. CTX in WT, KO vs. WT in CTX, and KO vs. WT in IM with log_2_FC of accessibility in the promoter-TSS region (DARs; FDR ≤ 0.05). Stained in red and blue are significant DEGs with significant DARs in the promoter-TSS region (|log_2_FC| ≥ 1, FDR ≤ 0.05). Pearson correlation coefficient r_sig_ was calculated from significant DEGs with DARs in promoter-TSS. Shaded in grey are not significant DEGs with DARs with |log_2_FC| ≤ 1. Pearson correlation r_all_ was calculated from significant (coloured) and not significant (grey) DEGs with DARs in promoter-TSS. (c) Coverage tracks of RNA and ATAC peak signals in WT and NFAT5-KO lines of CTX and IM at genes with higher expression in CTX (left) and at genes with higher expression in the IM (right).

In the context of comparative analysis between IM-WT and CTX-WT, a subset of up-regulated genes including *Nfat5* and *Aqp2*, showed a positive correlation with gain in promoter accessibility (up/gain) within 18.5% (685 out of 3,705) of these DEGs. Conversely, within the content of genes with decreased expression in the IM-WT, a proportion of 37.1% (1,101 out of 2,968) exhibited a positive correlation between loss in chromatin accessibility and decreased expression (down/loss). Only a minority subset of DEGs displayed unexpected transcriptional expression patterns relative to their chromatin accessibility changes. Specifically, only 4% (148 out of 3,705) of DEGs exhibited up-regulation in spite of loss in accessibility (up/loss), while 4.4% (131 out of 2,968) showed down-regulation in spite of gain in accessibility (down/gain). In the CTX-KO, a distinct subset comprising 5.8% (110 out of 1,911) of up-regulated DEGs due to the absence of NFAT5, demonstrated increased promoter accessibility (up/gain). This is in contrast to a smaller subset of 4.8% (10 out of 207) of DEGs showing both decreased expression and accessibility (down/loss).

Notably, within the CTX-KO, no genes were identified where up-regulated gene expression occurred simultaneously with closed chromatin. Conversely, only a single DEG showed down-regulation despite open chromatin conformation. Furthermore, there was a distinct difference between IM and CTX, especially with regard to the effects of NFAT5 depletion on the different gene expression landscape. In the IM-KO, a larger proportion of up-regulated DEGs (7.9%; 215 out of 2,720) exhibited a positive correlation with a gain in promoter accessibility, whereas 17.4% (79 of 454) of down-regulated DEGs showed a positive correlation with more closed promoter regions. In contrast, in the IM-KO, only a small proportion of 0.09% DEGs (24 out of 2,720) demonstrated increased transcriptional activity in spite of a closed chromatin environment. Moreover, a single DEG (1 out of 454) displayed down-regulation despite an observed increase in accessibility (Fig. 4a).

In the analysis on Figure 4a and b we focused on DARs that are localized at the promotor-TSS. We are ware, that enhancer regions, that could modulate gene expression level could be localized several kb up-or downstream of the TSS. We have therefore compared the accessibility of all annotated gene regions (promoter-TSS, exon, intron, TTS and intergenic) in IM with gene expression. This analysis showed that for 76.5% of all up-regulated DEGs correlate with higher chromatin accessibility and 72.8% of down-regulated DEGs with closed chromatin (up/gain and down/loss; Supp. Fig. 1, supplementary data S5).

Furthermore, Pearson correlation analyses obtained in CTX and IM of WT and NFAT5-KO demonstrate the positive correlation (r_sig_ ≥ 0.7) between significant alterations in promoter accessibility and changes in gene expression for the DEGs, including *Nfat5* and *Aqp2* (Fig. 4b). As an example, Figure 4c visualizes the tracks from RNA-seq and ATAC-seq for six genes that show differentially highest gene expression in either CTX or IM. The chromatin accessibility and transcription level of *Slc5a12* (solute carrier family 5, member 12), *Enpp6* (ectonucleotide pyrophosphatase/phosphodiesterase 6) and *Slc12a3* (solute carrier family 12, member 3) in CTX were higher than those in IM. On the contrary, the chromatin accessibility and transcription level of *Aldh1a3* (aldehyde dehydrogenase family 1, subfamily A3), *Wnt4* (wingless-type MMTV integration site family, member 4) and *Upk3bl* (uroplakin 3B-like) in IM were higher than those in the CTX.

The observed differences in gene expression are associated with correlating changes in promoter region accessibility and offer a possible explanation for the spatial differences in gene expression. These results highlight the relationship between gene expression and chromatin accessibility in the context of spatial localization and NFAT5-promoted regulation. The observed patterns suggest that NFAT5 plays a central role in modulating chromatin accessibility, and thereby influences gene expression in the IM and CTX regions of the kidney.

### Chromatin accessibility and gene expression correlates with spatial localization along the osmotic gradient

The functional diversity of the kidney segments is related to about 20 different highly specialized cell types of nephron and collecting duct (Fig. 5a)^45^. To analyze the correlation of gene expression and chromatin accessibility in the kidney we used public expression data from single cell analysis (https://cello.shinyapps.io/kidneycellexplorer/) ^46^ and mapped the data of genes with the most differentially accessible promoter regions between CTX and IM (Fig. 5). The top ten genes with gain or loss of promoter accessibility between CTX and IM show a specific spatial expression pattern (Fig. 5b and c). As expected, genes with gain of promoter accessibility in CTX have the highest expression in the different cell types that are located in CTX (proximal tubule; Fig. 5b), while genes with gain in promoter accessibility in IM show enriched expression in cells that are located in the IM, especially in cells of the inner medullary collecting duct (Fig. 5c). In this area, the renal cells are exposed to a hypertonic environment. The observed correlation between chromatin accessibility and gene expression with spatial localization along the osmotic gradient, supporting the hypothesis that hypertonicity and the corticomedullary osmotic gradient could probably be the main factors affecting the chromatin accessibility and renal specific gene expression pattern.

**Figure 5:**
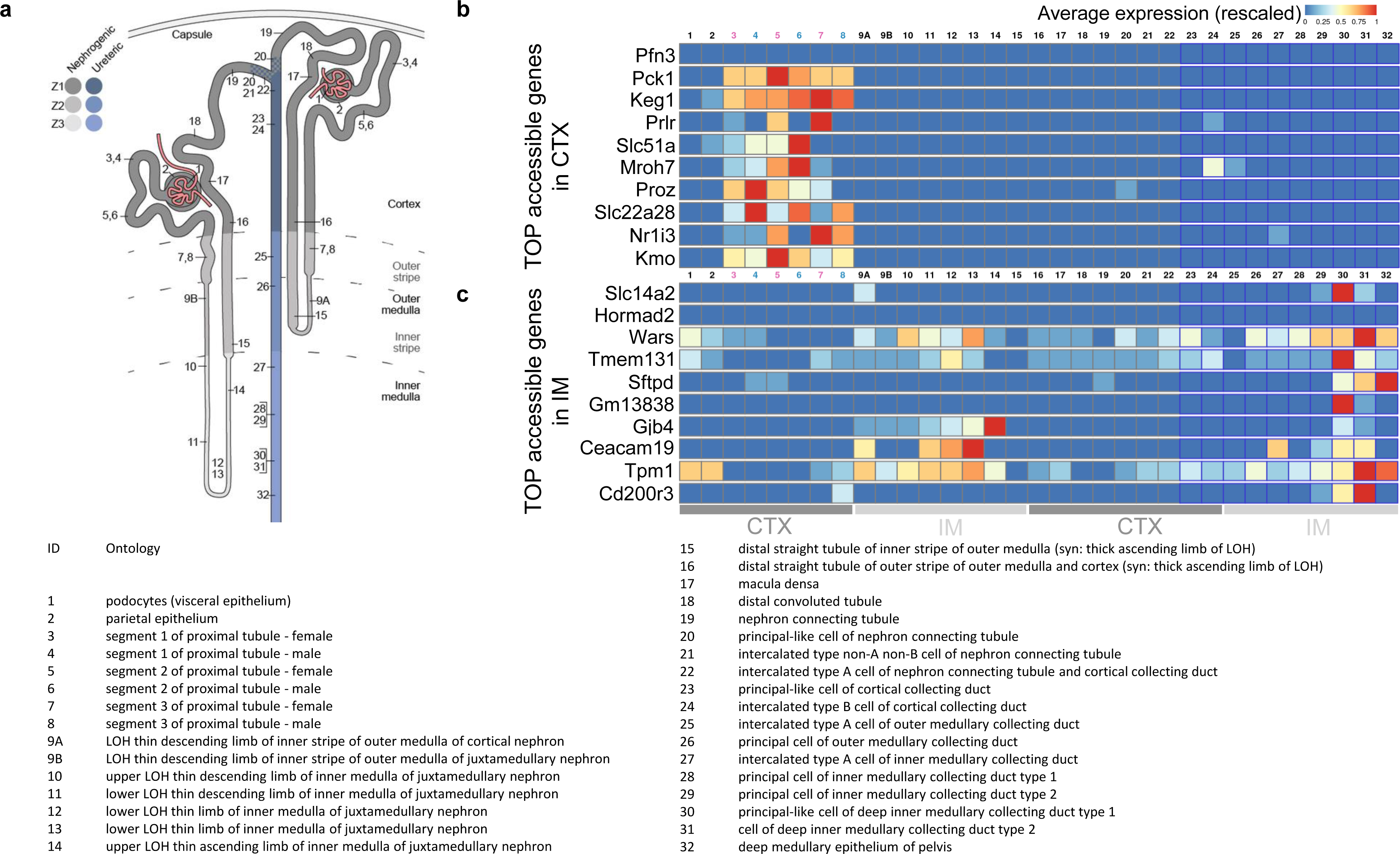
Correlation of chromatin accessibility data and public spatial gene expression data along the osmotic gradient. (a) Schematic map indicating the anatomical location of numbered metacells of the nephron and collecting duct in heatmap b and c. The ontology terms of numbered metacells are described in the legend below. The Kidney Cell Explorer (https://cello.shinyapps.io/kidneycellexplorer/; ^46^) was used to visualize the heatmap of spatial gene expression patterns of ATAC-seq resulting genes with the most accessible promoter (−5 kb −1 kb from transcription start site) of wild-type CTX (b) and IM (c) in the different kidney metacells. The color bar represents the rescaled average expression.

### Spatial localization and loss of NFAT5 affects accessibility at binding sites for other transcription factors

Changes in chromatin accessibility have profound effects on the binding of transcription factors (TFs) to their corresponding binding motifs and finally influence transcriptional activity and gene expression. Therefore, we performed DNA motif enrichment analysis to identify enriched TF motifs within the DARs and focused on those that positively correlated with DEGs (Fig. 4b). Our results revealed distinct differences in the enrichment of binding motifs between CTX and IM highlighting their changes in chromatin accessibility and gene expression (Fig. 6, supplementary data S6).

**Figure 6:**
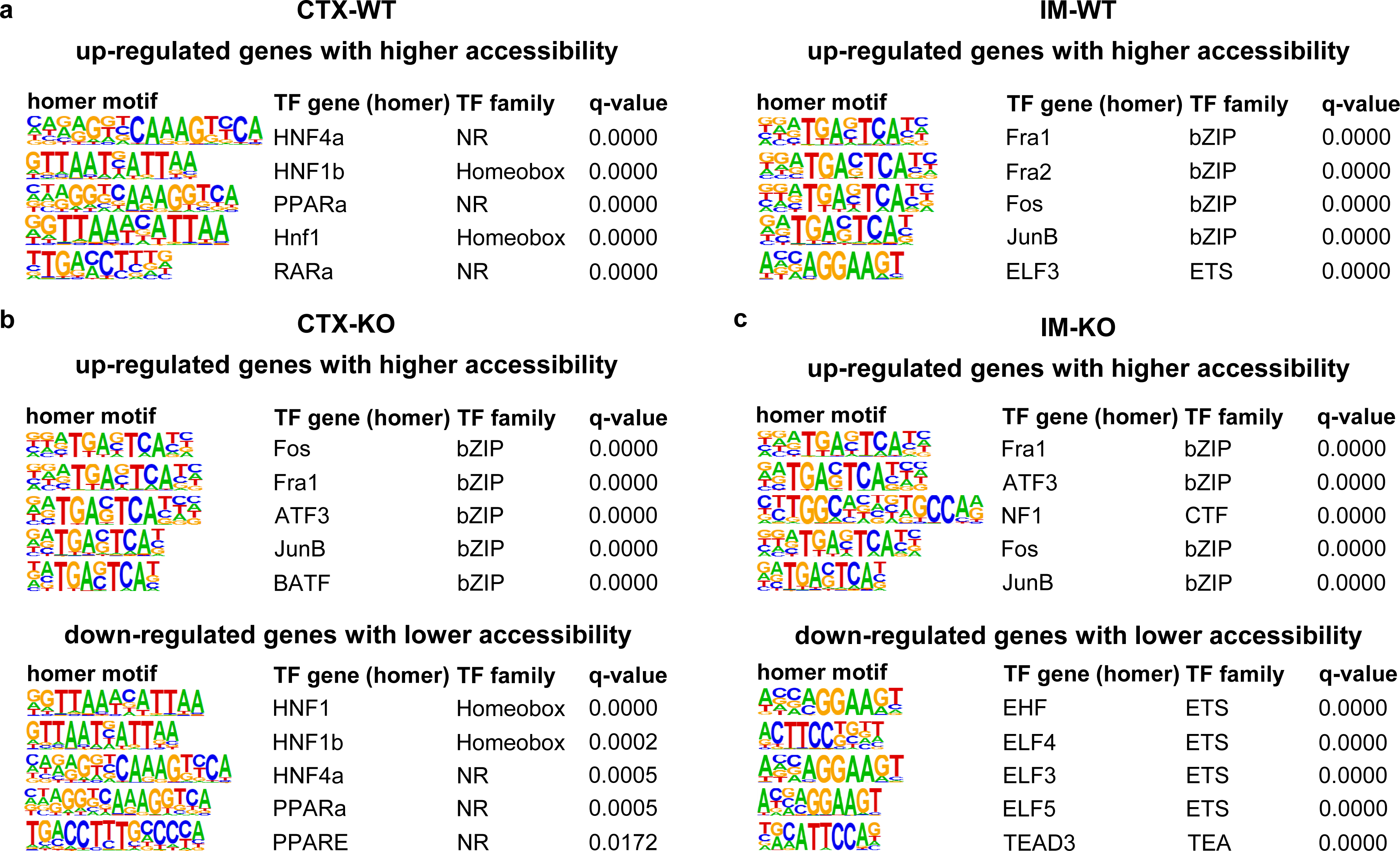
Changes in spatial localization and loss of NFAT5 alter the accessibility of transcription factor binding sites. (a) Enriched transcription factor (TF) motifs in regions with higher accessibility of increased expressed genes comparing CTX-WT and IM-WT. Enriched TF motifs for CTX (b) and IM (c) in the more accessible regions of higher expressed genes (top) and less accessible regions of lower expressed genes (bottom) comparing NFAT5-KO vs. WT. The complete lists can be found in supplementary data S5.

In the CTX, a total of 244 TF binding motifs showed significant enrichment (q-value <0.001) in DEGs with higher expression and increased accessibility. Similarly, in the IM, 196 motifs were enriched. Among these, 99 motifs were common in both WT CTX and IM, while 145 motifs were exclusively present in the CTX and 97 motifs were unique to the IM.

Comparing analysis between control CTX and IM, revealed a strong enrichment of motifs associated with the nuclear receptor (NR) family, such as HNF4α, PPARα and RARα, along with the homeobox family, including HNF1/HNF1α, in the CTX. In contrast, the IM-WT exhibited enriched binding motifs mainly for the βZIP family (including Fra1/Fra2, JUNB/FOS) and for the ETS family with ELF3 (Fig. 6a). To our surprise, NFAT5 motifs were not enriched in the IM.

The loss of NFAT5 resulted in a similar effect in both the CTX and IM, leading to increased accessibility of βZIP family motifs (Fig. 6b, upper illustration). These βZIP motifs were already accessible in the IM-WT and their accessibility was not affected by the loss of NFAT5. In contrast, in the CTX, the absence of NFAT5 led to higher accessibility of IM-WT specific motifs. Remarkably, the loss of NFAT5 increased the accessibility of 65 motifs in CTX and 102 motifs in IM among higher expressed DEGs. Notably, 16 motifs were uniquely accessible in the CTX, and 53 motifs were unique to the IM.

Furthermore, the study identified differences in binding motifs between the CTX-KO and IM-KO in less accessible regions associated with reduced gene expression. Strikingly, NR and homeobox family binding motifs specific to CTX-WT exhibited reduced accessibility in the CTX-KO due to loss of NFAT5 (Fig. 6b, lower illustration). In the IM-KO, chromatin packaging of ETS transcription factor motifs (bound by EHF, ELF3/4/5) was noticed, along with decreased expression of motif-associated genes (Fig. 6c, lower illustration). Interestingly, limited overlap was found upon analysis of the less accessible motifs in down-regulated DEGs, with only one common motif identified in both CTX and IM, while three motifs were unique to the CTX and 39 motifs were unique to the IM.

These findings highlight the function of NFAT5 in shaping tissue-specific transcriptional landscapes by influencing the binding motifs and accessibility of transcription factors. Overall, these results give important insights into the interaction between chromatin accessibility, TF binding motifs, and gene expression and identify the specific TF families related to different regions within the kidney.

### The spatial localization and loss of NFAT5 affect the accessibility of genes associated with the metabolic network, intracellular transport, cellular stress responses, and inflammatory and immune responses

Gene ontology (GO) biology process annotations of the genes with more accessible promoters in the CTX-WT revealed a significant enrichment of genes related to catabolic and metabolic processes such as catabolic processes of organic substances and macromolecules and metabolic processes of cellular amides and peptides, but also genes related to cellular response to stress and intracellular transport (Fig. 7a). In contrast, IM genes with more accessible promoters are involved in developmental and morphogenetic processes, such as tissue and epithelium development and morphogenesis and development of the tubes. In addition, these genes are associated with homeostasis and regulation of transport (Fig. 7a). The loss of NFAT5 in CTX mainly affect a gain of promotor accessibility of genes which play roles in inflammatory, immune and defense responses (Fig. 7b, upper illustration) while in the IM more accessible genes participate in positive regulation of cell communication, signaling and transport processes of ion and cations (Fig. 7c, upper illustration). Genes in CTX and IM, whose promoters are less accessible because of the loss of NFAT5, are mainly involved in the cellular response to stress, intracellular transport and catabolic processes of macromolecules, proteins and organic substances (Fig. 7b and c, lower illustrations). Most importantly, the reduced accessibility of stress response genes fits with the function of NFAT5 as an important regulator of adaptation to osmotic stress.

**Figure 7:**
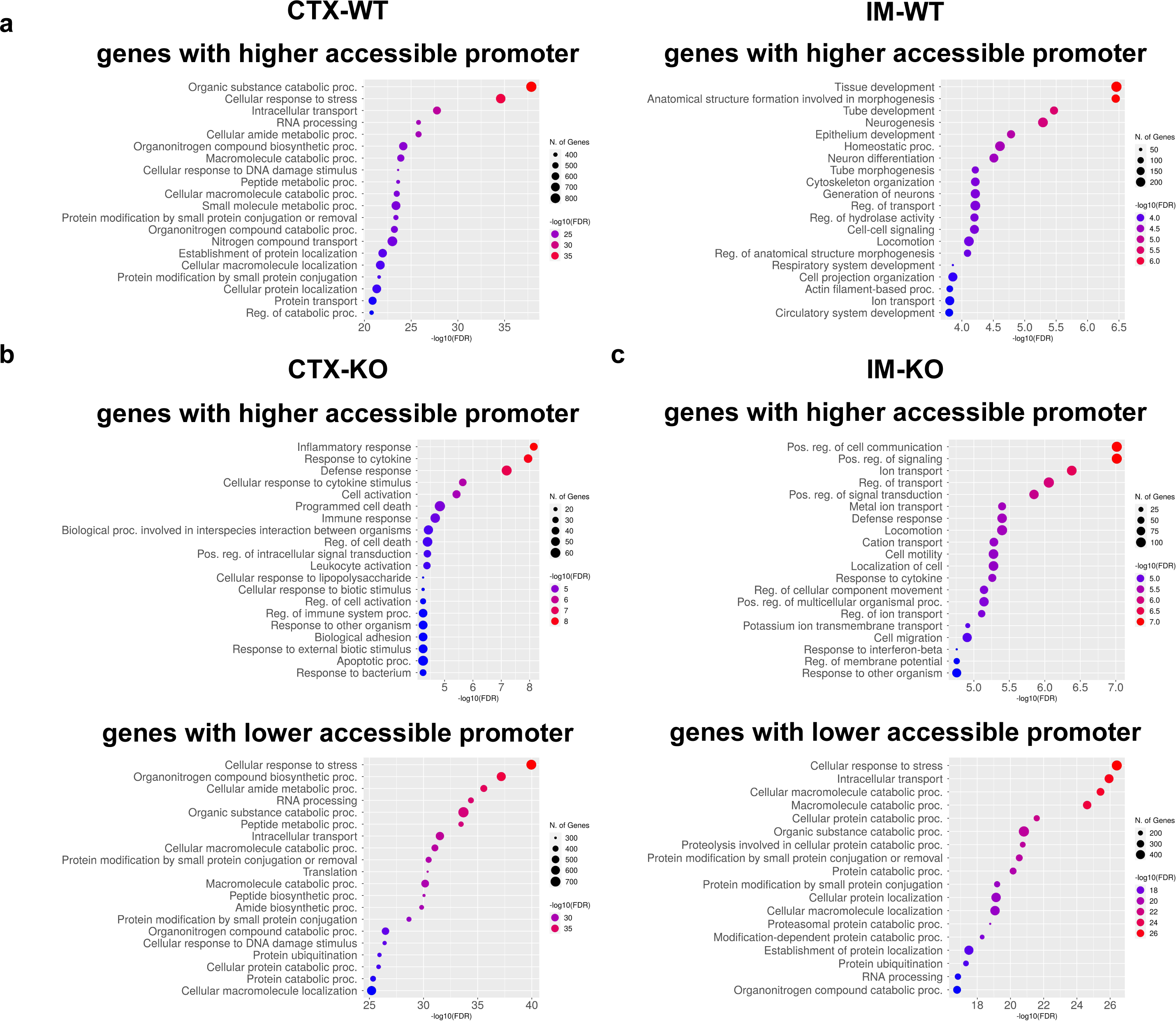
Spatial localization and NFAT5 absence affects the accessibility of genes involved in several biological processes. (a) TOP20 of Gene Ontology (GO) biological processes (FDR ≤ 0.05) with genes having a higher accessible promoter in CTX-WT and IM-WT (log_2_FC ≥ 0, FDR ≤ 0.01). TOP20 of GO biological processes (FDR ≤ 0.05) with genes having a higher or lower accessible promoter in CTX (b) and IM (c) after loss of NFAT5 (|log_2_FC| ≥ 0, FDR ≤ 0.01).

### Higher expression of *Aqp2* in mouse renal IM and in human collecting duct cells is promoted by increased chromatin accessibility

There are major differences in the gene expression pattern between renal CTX and IM. The corticomedullary osmotic gradient is probably one of the main factors influencing the gene expression profile. Our data show that chromatin accessibility and the effect of NFAT5 play an important role in influencing the gene expression profile. The NFAT5-KO mice exhibit a nephrogenic diabetes insipidus-like phenotype that is likely due to decreased expression of AQP2 ^32^. In primary cultured IMCD cells, the expression of *Aqp2* is strongly induced under hypertonic cell culture conditions ^28^. Furthermore, single-cell RNA sequencing from mouse kidney showed that the renal gene expression of *Aqp2* is highest in IM ^46^.

We therefore investigated whether the expression of *Aqp2* is regulated by differences in chromatin accessibility. And indeed, higher *Aqp2* mRNA expression in the IM correlates significantly with the higher accessibility of the *Aqp2* promoter region in the IM compared to CTX (Fig. 4b left, Fig. 8a). This suggests that hypertonicity leads to an improvement in the accessibility of the promoter region of *Aqp2*. In IM-KO, and also in CTX-KO, the accessibility of the promoter was decreased, which could explain the reduced *Aqp2* expression in NFAT5-KO mice (Fig. 4b, Fig. 8a). The accessibility of this promoter region was also identified in the developing mice kidney (Fig. 8b), indicating an important function of this region ^45^.

**Figure 8:**
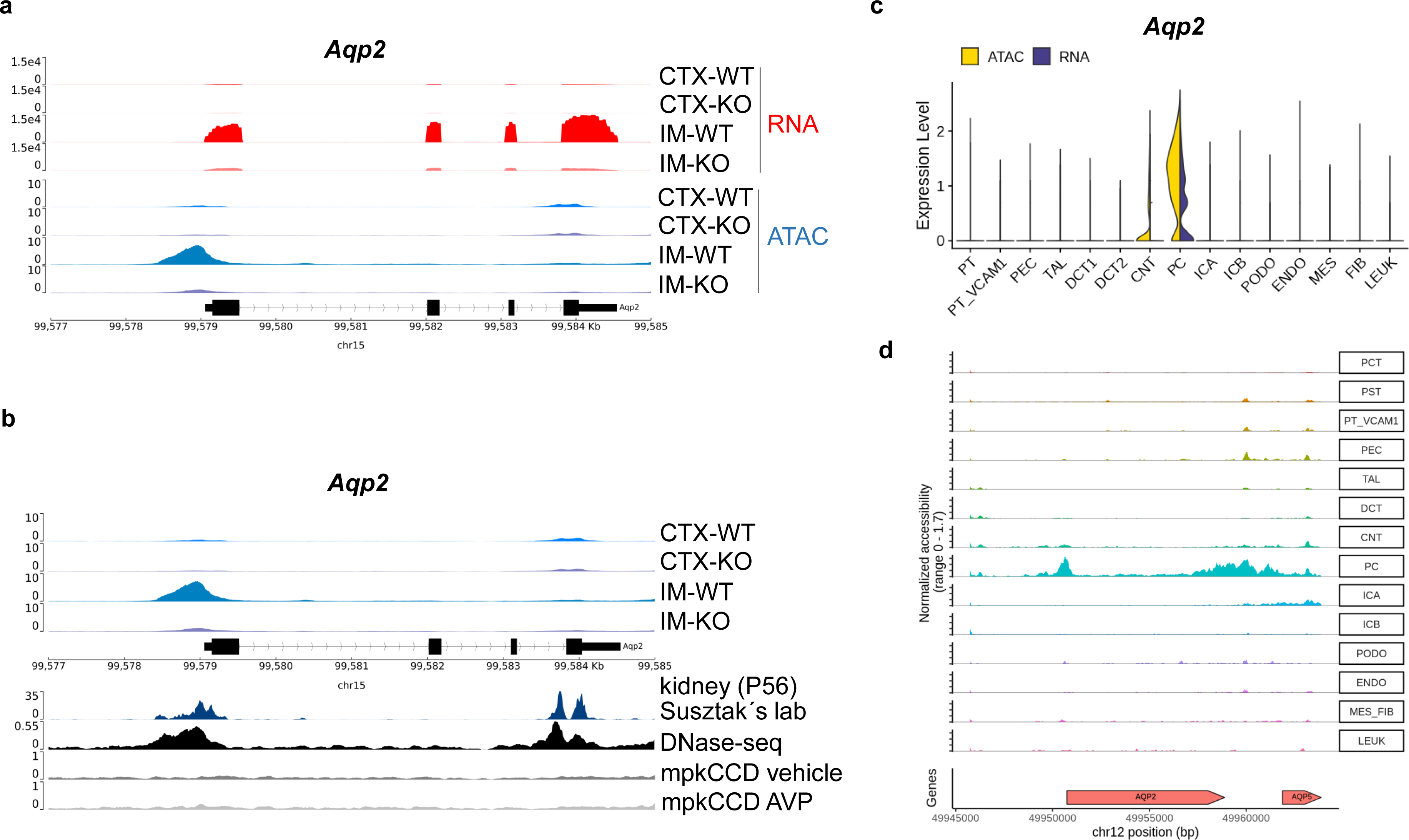
The enriched expression of *Aqp2* is promoted by increased accessibility of the promoter region. (a) Coverage tracks of RNA and ATAC peak signals in WT and NFAT5-KO lines of CTX and IM for *Aquaporin 2 (Aqp2)*. (b) Coverage tracks of ATAC peak signals in WT and NFAT5-KO lines from CTX and IM for *Aqp2*. To compare the CTX and IM results with public ATAC-kidney and ATAC-mpkCCD (murine principal kidney cortical collecting duct) cells open access ATAC data for developing kidney (P56-Susztak’s lab, https://esbl.nhlbi.nih.gov/IGV_mo/ ^45^), DNase-seq samples from adult mice kidney (GEO:GSM1014193 ^45^) and mpkCCD cells treated with vehicle or vasopressin (AVP, GSE108786 ^47^) were added. (c) ATAC and RNA expression level of *Aqp2* from single-cell data of major cell types in the adult human kidney: proximal tubule (PT), parietal epithelial cells (PEC), thick ascending limb (TAL), distal tubule (DCT1, DCT2), connecting tubule (CNT), collecting duct (PC, ICA, ICB), endothelial cells (ENDO), glomerular cell types (MES, PODO), fibroblasts (FIB), and a small population of leukocytes (LEUK). (d) Single cell ATAC tracks of major human kidney cell types mentioned in (c) for *Aqp2*. Human single cell data from (c) and (d) are from Muto et al., 2021 ^1^ and were visualized using the KIT platform (http://humphreyslab.com/SingleCell/).

AQP2 is an important marker for the major cells of the collecting duct. The collecting duct cells are localized in the CTX of the kidney, in the IM and even in the renal papilla and are exposed to a hypertonic environment similar to the cells localized in the IM. The expression of *Aqp2* is also induced by continuous stimulation with vasopressin (AVP). To compare our CTX and IM ATAC results with mpkCCD cells, open-access ATAC track data for these cells treated with vehicle or AVP ^47^ were examined for the *Aqp2* gene. No improvement in accessibility was observed in the promoter region of *Aqp2* by AVP (Fig. 8b). However, similar to the study with the developing kidney, this region was also present in DNase treated adult mice (GEO Acc. No. GEO:GSM1014193 ^45^). Open-access single-cell data from the major cell types in the adult kidney also demonstrate in humans that a high *AQP2* expression in the principal cells (PC) of the collecting duct correlates with the accessibility of *AQP2* (Fig. 8c). Similar to the situation in mice, accessibility gain in these human PC cells is limited to the promoter region of *AQP2* (Fig. 8d), implicating a conserved control of transcriptional regulation.

## DISCUSSION

Cell specific gene expression is often associated with epigenetic modifications including DNA methylation, histone modification or chromatin accessibility. Using the Assay for Transposase-Accessible Chromatin (ATAC-seq), it is possible to identify open chromatin regions. These kind of changes have been analyzed in using kidney samples to identify physiological or pathophysiological in DNA methylation, histone modification or chromatin accessibility ^48^. Meanwhile several studies have been performed using single cells, renal tissues or nephron segments to correlate gene expression with differences in chromatin accessibility using ATAC-seq ^49–51^. With these and other studies, it was possible to identify a nephron segment or cell type specific gene expression pattern that might be associated with epigenetic modifications. The factors that control these changes are not fully understood. There is plenty of evidence that the unique hypertonic environment in the renal inner medulla mediates a specific gene expression pattern ^12, 29, 52^ ^31^. With this study, we provide evidences that the differences in the chromatin accessibility contribute to the observed changes.

Between mouse renal cortex and renal inner medulla, more than 10,000 genes are differentially expressed ^29^. The analysis of genomic regions showed that more than 60,000 regions are commonly accessible in renal CTX and renal IM. Nearly the same number of regions (41,979) are uniquely accessible only in the CTX and around 48,000 regions only in the IM. The number of unique accessible regions also reflect the number of DARs between CTX and IM. When we mapped the genes that are associated with the top ten DARs between renal CTX and IM using the kidney single cell explorer ^7^, we showed that the genes with higher accessible regions in CTX are nearly exclusively expressed in cells of the proximal tubule. In the same way, genes that have higher accessibility in the IM have the highest expression in cells, which are located in the IM. The cells in the IM might be influenced by a hypertonic environment, suggesting that differences in osmolality could probably regulate gene expression by modulating chromatin accessibility.

The correlation of DEGs with DARs also shows that the differences in gene expression between CTX and IM is associated with changes in the accessibility in the TSS of the genes. We are aware that the enhancers can also be located several kb upstream and downstream of the TSS and modulate the expression level. When we compare the DEGs with all DARs we get even a higher correlation of DEGs with DARs (supplementary figure S1).

When we performed transcription factor motif analysis that could bind to the open chromatin regions, we observed differences between CTX and IM. In the CTX for example motif analysis identified enriched motifs for the TFs Hnf1b and Hnf4a in the DARs. Hnf1b and Hnf4a are proximal tubulus specific TFs ^53, 54^. Our data indicates that the action of these TFs is associated with increased accessibility at the TSS of the target genes. Within the top five TFs in the IM for JunB and Elf3, studies have shown a possible involvement in the regulation of *Aqp2* expression ^55–58^. *Aqp2* has its highest expression level in the IM ^2, 7, 29, 52^. Our data show, that a contribution of these factors is supported by increased accessibility of the motifs for these TFs at the TSS of *Aqp2*. The transcriptional regulation of *Aqp2* is mediated by the action of the antidiuretic hormone vasopressin (AVP) ^59^. Using mpkCCD cells, an AVP sensitive collecting duct cell line, for ATAC-seq and integration of ChIP-seq led to the identification of two enhancer regions, that are accessible in cells incubated with AVP ^60^. Potential enhancer regions were identified located 81 kb upstream and 5.8 kb downstream of *Aqp2* TSS. However, a region close to the TSS was not evident in AVP treated cells. Within this study, we have identified a region close to the TSS of *Aqp2* that shows higher accessibility in IM compared to CTX, which could explain the differences in the expression level of *Aqp2*. This region can also be found in other studies, where ATAC-seq have been performed using mice or human kidney samples ^45, 49, 61–63^. In these studies, a differentiation between CTX and IM is missing. The higher accessibility in IM correlates with expression of *Aqp2* and the hyperosmolality. This data indicates that the enriched *Aqp2* expression in the IM is associated with increased accessibility at the TSS and that probably hypertonicity induces the observed differences. Recently, Haug et al. analysed chromatin accessibility and spatial transcription in the adult human kidney cortex and medulla ^31^. Haug et al. identified around 2,300 differentially expressed genes between renal cortex and renal medulla. Using mice renal cortex and renal inner medulla, we were able to identify around 10,000 differentially expressed genes. This is probably due to the fact, that we used renal inner medulla, including renal papilla, for analysis. In this region, the environmental hypertonicity is higher and this influences also the gene expression level ^52^.

The environmental hypertonicity induces the activation of NFAT5 that activates an osmoprotective gene expression profile ^24, 25, 64^. Surprisingly the analysis of TF motifs did not show a significant enrichment of NFAT5 motifs in the IM for DAR regions. Loss of NFAT5 in the induced a diabetes insipidus like phenotype and this was associated with massive changes in gene expression ^29, 32^. The changes in gene expression are also associated with differences in chromatin accessibility. Loss of NFAT5 led to identification of around 34,000 novel accessible genomic regions. From these around 8,700 are unique to CTX and 19,700 to IM. This is also reflected by the analysis of DARs between control and NFAT5 CTX and IM. The analysis of the TF motifs showed that loss of NFAT5 in the cortex lower accessibility is associated with Hnf1b, Hnf4a or PPARa, all factors that showed enrichment in control CTX. In the IM reduced gene expression and accessibility was associated with EHF, ELF3 or ELF5 TF motifs. ELF5 is a principal cell specific TF and might be involved in *Aqp2* expression ^55^. However, a collecting duct principal cell specific ELF5 deficient mice showed only minor differences in *Aqp2* expression ^32^.

Several studies using cell lines and native tissue have shown, that hypertonicity and NFAT5 are master regulators of *Aqp2* expression ^29, 32, 52, 65^. Here we showed that the increased *Aqp2* expression in the IM is associated with a gain of accessibility of *Aqp2* at the promotor region. In NFAT5 deficient IM the accessibility is massively decreased and correlates with reduced *Aqp2* mRNA expression, highlighting the role of NFAT5 in *Aqp2* expression. Based from the data we can only speculate, if NFAT5 directly affects the accessibility in the *Aqp2* promotor region or another NFAT5 target gene is involved in the observed changes.

The loss of NFAT5 is linked to renal inflammation and an up-regulation of kidney injury marker genes ^29^. The underlying mechanism for this increased expression of immune-related genes is likely due to an increased infiltration of immune cells, indicated by the elevated expression of *Cd4*, *Cd8,* or *Cd14* ^29^. Higher gene expression depends on the condition of enhanced accessibility of these genes. In relation to the loss of NFAT5 in the CTX specifically, there is evidence of gain in promoter accessibility for genes associated with inflammatory, immune, and defense responses (Fig. 7). This complex network of immune cells infiltration and regulatory processes offers a plausible explanation for the observed increased expression of immune-related genes in NFAT5-KO mice. NFAT5 may influence the promoter accessibility and modify the dynamics of the immune in the kidneys.

In conclusion, our study, using ATAC-seq, provides insights into the global differences in chromatin accessibility and gene expression patterns between renal CTX and IM. The results highlight the role of spatial localization and NFAT5 in modulating chromatin region accessibility, particularly at promoters, and their correlation with changes in renal gene expression. Understanding these epigenetic mechanisms in the kidney can enhance our knowledge of renal physiology and pathophysiology, particularly in the context of osmotic gradients and renal diseases. These findings highlight the importance of epigenetic regulation in shaping spatial gene expression patterns in the kidney and provide a basis for further investigations into the molecular mechanisms underlying renal function and dysfunction.

## DATA STATEMENT

The ATAC-seq raw fastq files and bigwig files will be deposited in the repository Gene Expression Omnibus with the accession number GSE247097, https://www.ncbi.nlm.nih.gov/geo/query/acc.cgi?acc=GSE247097). The RNA-seq data are available under the accession code GSE195881, https://www.ncbi.nlm.nih.gov/geo/query/acc.cgi?acc=GSE195881) ^29^. We have clearly indicated each software and provided information on the parameters selected (METHODS). The authors declare that all other data supporting the conclusions of this article are included within the article and its supplementary files.

## ACKNOWLEDGEMENTS

The work was funded by the Deutsche Forschungsgemeinschaft (DFG, German Research Foundation) –ED181/9-3. The funder had no role in study design, data collection and analysis, decision to publish, or preparation of the manuscript.

## AUTHOR CONTRIBUTIONS

K.E., D.C. and K.N. performed the experiments, K.S., A.D. and K.E. analyzed the data, B.E. supvervised the project, K.E. and B.E. wrote the manuscript, B.E. designed the study. All authors read and approved the final manuscript.

## SUPPLEMENTARY MATERIAL

**Supplementary figure S1**. Correlation of gene expression and chromatin accessibility. (A) Comparison of differentially expressed genes (DEGs; FDR ≤ 0.05, |log_2_FC| ≥ 1) with differentially accessible regions (DARs; FDR ≤ 0.05, |log_2_FC| ≥ 1). Shown is the proportion of genes with up-regulated gene expression and gain-or loss of chromatin accessibility (up/gain and up/loss) and with down-regulated gene expression and gain-or loss of chromatin accessibility (down/gain and down/loss) in the four gene sets of the CTX and IM of WT and NFAT5-KO mice. The corresponding genes are listed in supplemental data S4.

**Supplementary data S1.** List of accessible regions in renal cortex and inner medulla of WT and NFAT5-KO mice.

**Supplementary data S2.** List of consensus peaks set.

**Supplementary data S3.** List of differentially accessible regions between renal cortex and inner medulla of WT and NFAT5-KO mice (DESEQ2 results) from Fig.3.

**Supplementary data S4.** Venn diagrams and lists of genes from intersection of RNA-seq and promoter-TSS ATAC-seq data from Fig.4a.

**Supplementary data S5.** Venn diagrams and lists of genes from intersection of RNA-seq and ATAC-seq data

**Supplementary data S6.** Lists of known motifs from TF motif enrichment analysis from Fig.6.

**Supplementary references.** Chernyakov, D., et al., The nuclear factor of activated T cells 5 (NFAT5) contributes to the renal corticomedullary differences in gene expression. Sci Rep, 2022. 12(1): p. 20304.

